# Mechanistic origins and evolutionary erosion of collateral sensitivity in a β-lactamase

**DOI:** 10.64898/2026.02.12.705688

**Authors:** Daniel Salamonsen, Karol Buda, Daojiong Wang, Kinga Virág Gulyás, Marc W. van der Kamp, Christopher Frøhlich

**Author notes:** Corresponding author: Christopher Frøhlich.

## Abstract

As antibiotic discovery stalls, exploiting collateral sensitivity, where resistance to one drug increases sensitivity to another, offers a promising route to extend the lifespan of existing drugs. However, the molecular origins and robustness of such trade-offs at the level of single resistance determinants remain poorly understood. Here, we examined a previously evolved trajectory of the β-lactamase OXA-48 to Q4 (A33V/F72L/T212A/S213A), which confers a 40-fold increase in ceftazidime resistance when expressed in *Escherichia coli*. This evolution coincided with the emergence of collateral sensitivity to piperacillin (27-fold reduction), likely caused by the introduction of F72L. We challenged the stability of this collateral sensitivity network by subjecting Q4 to directed evolution under co-selective pressure from both ceftazidime and piperacillin. This selected for the substitution V120G, which alleviated the piperacillin trade-off while maintaining elevated resistance to ceftazidime in genetic backgrounds harboring F72L. Structural and computational analyses revealed that, during evolution from OXA-48 to Q4, F72L introduced substantial conformational changes in the Ω-loop, likely leading to less productive piperacillin binding poses. V120G counteracted the effect of F72L by decreasing the Ω-loop’s conformational freedom, thereby partially restoring piperacillin resistance. Finally, we show that other substitutions at position 120 can exert similar mitigating effects. Taken together, our results provide a mechanistic understanding of how adaptive solutions both generate and erode collateral sensitivity, knowledge crucial for predicting the long-term stability of these networks.

**IMPORTANCE:** The evolution of antimicrobial resistance to one drug can also increase bacterial susceptibility to other drugs, a phenomenon known as collateral sensitivity. Understanding the mechanisms underlying these effects may help clinicians design antimicrobial treatment strategies that limit the emergence of resistance. However, for clinical applications, collateral effects must be stable and robust over a certain evolutionary time. While many studies have focused on the stability of collateral effects at the bacterial population level, their stability during the evolution of a single resistance determinant has remained largely unknown. Here, we studied the molecular origins of collateral sensitivity arising during the evolution of the β-lactamase OXA-48 and robustness of this network under co-selection. By linking resistance evolution to changes in protein structure, we illuminate the role of active site loop dynamics in shaping both the collateral effect and its mitigation. Such results are important for understanding collateral sensitivity–based strategies and how they may shape the evolutionary trajectories of resistance enzymes.

## INTRODUCTION

The impact of the global antimicrobial resistance crisis is amplified by a persistent innovation gap in the development of new antimicrobial agents. As the pipeline of novel drugs has been shrinking, extending the lifespan of existing antimicrobials and slowing the emergence of resistance have become an urgent priority^1^. One promising strategy involves exploiting the concept of collateral sensitivity, in which the evolution of resistance to one antimicrobial increases susceptibility to others^1–4^. Several strategies to take advantage of collateral networks have been proposed, including cycling-based regimens and combination therapy^5–15^. In particular, combining drugs that exhibit reciprocal collateral sensitivities can impose simultaneous selective pressures that restrict mutational escape, thereby maximizing bacterial killing and creating an evolutionary trap that limits adaptive pathways^8,14,15^.

The success of exploiting collateral sensitivity depends on multiple factors, including the evolutionary stability of these networks, which is shaped by genetic context, selection conditions, resistance mechanisms, and treatment design^1,3,10,16–20^. Experimental evolution studies have demonstrated that collateral responses can erode or invert, even in the absence of antimicrobial pressure^2,3,10,16,21,22^. However, these studies have mostly focused on the global response of bacterial populations. In contrast, collateral responses can originate from mutations in specific resistance determinants, such as β-lactamases^5,19,23–25^. Yet mechanistic understanding of how collateral sensitivity patterns arise and persist remains limited, particularly when a single resistance determinant evolves along defined evolutionary trajectories.

One such resistance determinant is the class D β-lactamase OXA-48, a major driver of antibiotic resistance globally due to its efficient hydrolysis of penicillins and carbapenems, as well as its plasmid-mediated dissemination^26^. The hydrolysis mechanism of OXA-48 relies on a catalytic serine (S70) and a carbamylated lysine (K73). Carbamylation of K73 is facilitated by the hydrophobic environment around the base, including the S118-V119-V120 motif, which lowers the local p*K*_a_ of K73^27^. Additionally, loop regions, such as the Ω-loop (Y144–R163) and the β5–β6 loop (T213–K218), influence substrate affinity and catalytic efficiency by shaping the active site^28^.

While OXA-48 efficiently hydrolyzes many penicillins and carbapenems (*k*_cat_/*K*_M_ up to 10^6^ M^−1^ s^−1^), it is a poor biocatalyst for the hydrolysis of 3^rd^ generation cephalosporins, particularly ceftazidime (CAZ), with a *k*_cat_/*K*_M_ around 10^2^ M^−1^ s^−1 29–31^. However, variants conferring increased CAZ resistance have been identified both in nature (e.g., OXA-163, OXA-247, OXA-405, and OXA-519) and during laboratory evolution (e.g., F72L), demonstrating that OXA-48 can readily adapt toward increased cephalosporin resistance^29,32–35^. We previously showed that acquisition of four mutations (Q4: A33V/F72L/S212A/T213) can enhance OXA-48-mediated CAZ resistance by ∼40-fold in *E. coli*. Importantly, these adaptations frequently coincide with a strong reduction in OXA-48’s ability to hydrolyze penicillins and carbapenems^32–35^. For example, the F72L substitution reduces the *k*_cat_/*K*_M_ for piperacillin (PIP) hydrolysis by 167-fold in OXA-48^29^.

Here, we provide a mechanistic study of the origin and stability of collateral sensitivity from an enzyme perspective, using the evolutionary trajectory from wild-type OXA-48 (wtOXA-48) to Q4^36^. First, we quantify how each mutation along the evolutionary trajectory affects the collateral relationship between CAZ and PIP resistance. Next, using directed evolution, we challenge the stability of the collateral network under dual selection with CAZ and PIP. By combining biochemical and structural analyses, we reveal how changes in the conformational flexibility of the Ω-loop are related to both collateral sensitivity against PIP and the erosion of this network.

## RESULTS

### Ceftazidime resistance evolution causes a strong trade-off with piperacillin

To investigate how collateral sensitivity develops along an evolutionary trajectory, we measured the half-maximal inhibitory concentrations (*IC*_50_) for piperacillin (PIP) in variants along the previously described CAZ trajectory (F72L → S212A → T213A → A33V, Fig. 1a-b, Tab. S1). F72L, which was the first acquired mutation, increased CAZ resistance by 2-fold while substantially reducing PIP resistance (23-fold) compared to wtOXA-48. Although CAZ resistance improved up to 40-fold along the evolutionary trajectory, PIP resistance remained consistently low after the introduction of F72L, resulting in a 27-fold reduction in *IC*_50_ compared to wtOXA-48. Notably, PIP resistance conferred by variants along the evolutionary trajectory remained >3-fold higher than that of the *E. coli* host strain lacking β-lactamase expression (Tab. S1).

**Fig. 1:**
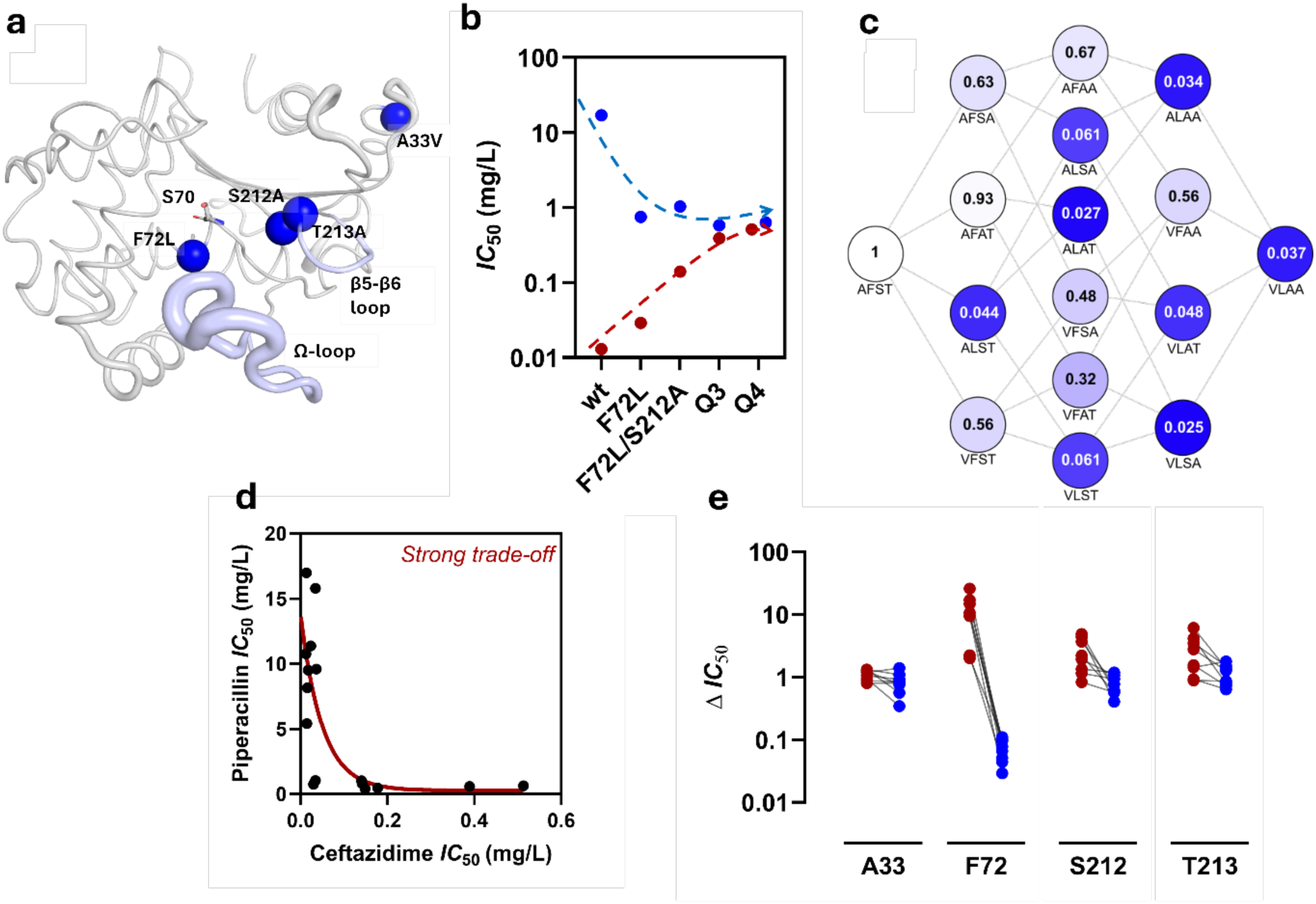
Trade-off development during evolution of ceftazidime resistance in OXA-48. **a.** Most of the acquired mutations are clustered around the active site, either in direct contact (F72L) or located in structural elements (S212A and T213A) relevant for substrate specificity (e.g., Ω-loop and β5-β6-loop). Created from PDB: 8PEB^36^. **b.** *IC_50_* measurements show that while ceftazidime resistance (red) progressively increased along the evolutionary trajectory, the acquisition of F72L introduced a substantial reduction in piperacillin resistance (blue) that persisted throughout subsequent evolution. **c.** The adaptive landscape of the four accumulated mutations in Q4 characterized by piperacillin resistance. Each node represents one of the 16 possible variants, labeled with single-letter amino acid codes, displaying the fold-change of the piperacillin *IC_50_* relative to wtOXA-48. Darker color indicates stronger collateral sensitivity in the form of reduced piperacillin *IC_50_*. **d.** Correlation between piperacillin and ceftazidime *IC_50_* values reveals a strong trade-off, whereby even minor improvements in ceftazidime resistance are associated with large reductions in piperacillin resistance. **e.** Difference in *IC_50_* fold-changes of piperacillin (blue) and ceftazidime (red) for the four accumulated mutations in all genetic backgrounds within the adaptive landscape, illustrating the dominant role of F72L in driving collateral sensitivity. Black lines connect the same genotype across substrates.

To quantify the contribution of individual mutations acquired along the evolutionary trajectory to the observed trade-off between CAZ and PIP, we determined *IC_50_* values across the adaptive landscape comprising all 16 mutational combinations (Fig. 1c, Tab. S1). Except for F72L, all other single mutants (A33V, S212A, and T213A) exhibited PIP resistance similar to wtOXA-48 (<2-fold change). Correlating *IC*_50_ values for both antibiotics revealed a highly concave collateral sensitivity relationship where mutations displaying even marginal improvements in CAZ resistance exhibited a strong deleterious effect on the conferred PIP resistance (Fig. 1d). Analyzing the *IC*_50_ fold-changes between PIP and CAZ across all genetic backgrounds revealed F72L as the primary driver for the PIP-CAZ trade-off (Fig. 1e). In summary, these results demonstrate that strong collateral sensitivity to PIP arises early in the evolutionary trajectory, is introduced by F72L, and remains stable even as CAZ resistance increases stepwise through subsequent mutations.

### Co-exposure selects for the trade-off-mitigating substitution V120G

We aimed to explore whether strong collateral effects can be mitigated under co-selective pressure from CAZ and PIP. To this end, error-prone PCR libraries were constructed from two evolutionary starting points: wtOXA-48, representing a variant that confers low CAZ and high PIP resistance (CAZ *MIC* 0.03 mg/L, PIP *MIC* 32 mg/L, Tab. S1), and Q4, which exhibits substantially increased CAZ resistance but a strong PIP trade-off relative to wtOXA-48 (CAZ *MIC* 1 mg/L, PIP *MIC* 2 mg/L, Tab. S1). The libraries were selected on agar plates containing increased concentrations of both CAZ and PIP simultaneously (Tab. S2).

No growth was detected from the wtOXA-48 library, even at the lowest tested antibiotic combination (CAZ 0.03 mg/L and PIP 4 mg/L). In contrast, selection of the Q4 library yielded three distinct variants at CAZ 0.15 mg/L and PIP 4 mg/L. *IC_50_* measurements showed that these variants increased PIP resistance by up to 5-fold relative to Q4 while largely retaining CAZ resistance (<2-fold reduction) (Tab. S3). All variants carried the V120G amino acid substitution along with one or two additional mutations, suggesting that V120G was likely under positive selection. Consistent with this, a reconstructed Q4:V120G variant maintained a 4-fold increase in PIP *IC_50,_* and a similar level of CAZ resistance compared to Q4 (Tab. S3).

Next, we asked whether the trade-off-mitigating effect of V120G depends on the specific mutations acquired during CAZ evolution. To address this, we extended our previous adaptive landscape by constructing all possible V120G combinations and determined their CAZ and PIP *IC_50_* (Fig. S1-S2, Tab. S4). Correlating CAZ and PIP *IC_50_* values for V120 and G120 variants revealed that G120 variants occupy intermediate positions between the two distinct clusters defined by F72- and L72-containing backgrounds, indicating a PIP trade-off–mitigating effect specifically in F72L-containing variants (Fig. 2a). In fact, V120G consistently increased PIP resistance in variants carrying F72L while being generally detrimental in backgrounds devoid of F72L (Fig. 2b). For instance, the introduction of V120G into wtOXA-48 led to a 14-fold reduction in PIP *IC*_50_, whereas it increased the PIP *IC*_50_ by 2-fold in combination with F72L. Similarly, the acquisition of V120G in the A33V/S212A/T213A triple mutant resulted in a 25-fold decrease in PIP *IC*_50_ while yielding a 4-fold increase in Q4. Conversely, V120G displayed only a marginal effect on CAZ resistance in all genetic backgrounds, with *IC_50_* fold changes reaching a maximum of 2-fold (Tab. S5).

**Fig. 2:**
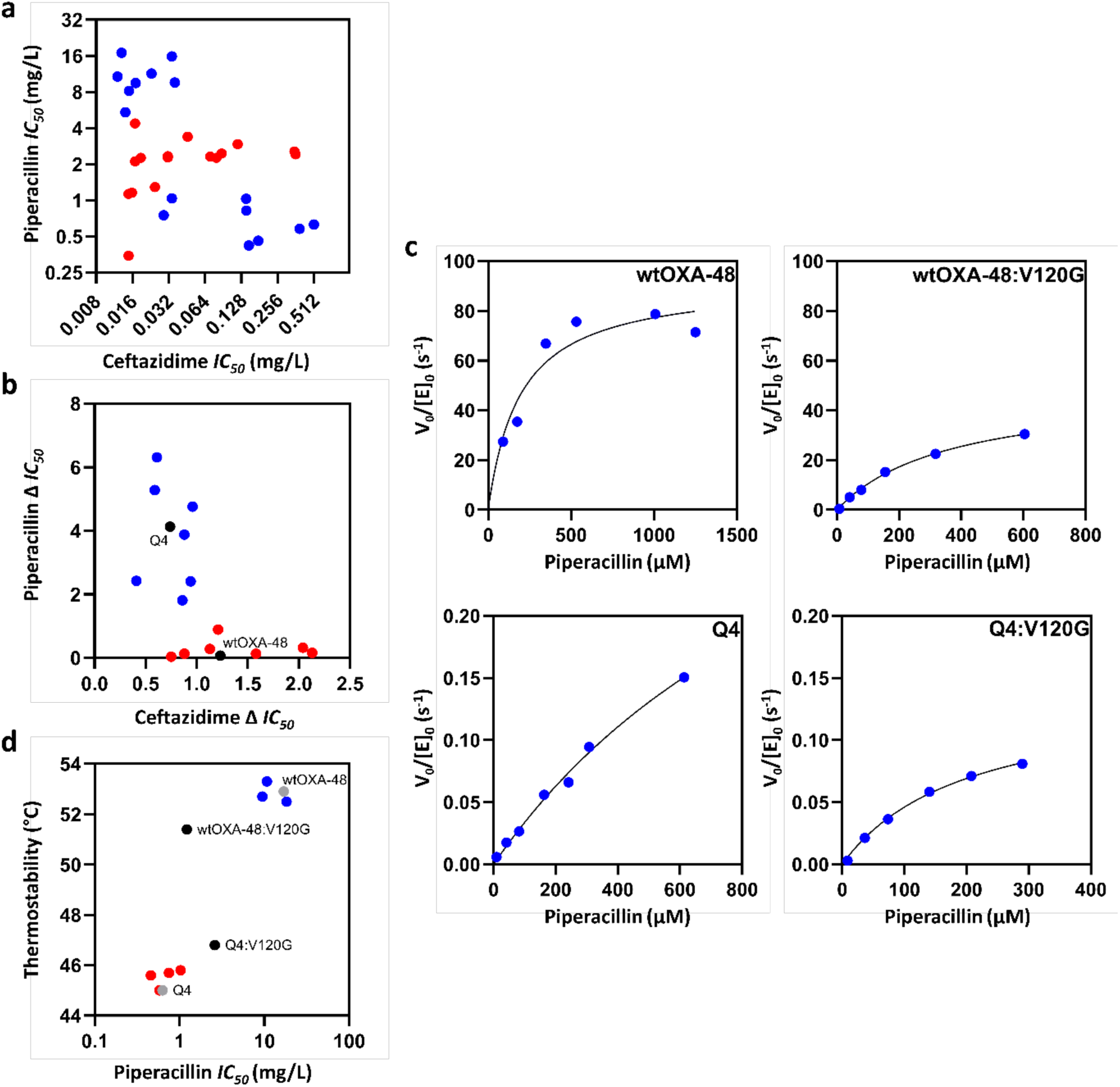
Collateral sensitivity and its erosion caused by the interplay of F72L and V120G. **a.** Landscapes of collateral sensitivity between ceftazidime and piperacillin *IC*_50_ for V120 variants (blue) and G120 variants (red), showing a mitigated trade-off in G120-containing variants (for all *IC*_50_ values and landscapes see Tab. S1 and Fig. S1-S2). **b.** *IC_50_* fold changes for ceftazidime and piperacillin resistance related to the addition of V120G (see Tab. S5). The mitigating effect of V120G on piperacillin resistance is highly dependent on the presence (blue) or absence (red) of F72L. wtOXA-48 and Q4 are highlighted in black. **c.** Steady-state kinetics for piperacillin hydrolysis by wtOXA-48, Q4, and their corresponding V120G variants. **d.** Thermostability (*T_M_*) *versus* piperacillin *IC*_50_ for selected variants, with wtOXA-48 and Q4 shown in grey and their V120G derivatives in black. Additional variants carrying (red) or lacking F72L (blue) from a previous study^36^ are included to illustrate the destabilizing effect of F72L.

To gain a mechanistic understanding of the context-dependent effect of V120G on PIP resistance, we performed steady-state kinetic analyses for wtOXA-48, Q4, and their corresponding V120G variants (Fig. 2c, Tab. 1). Q4 displayed a 325-fold reduction in PIP turnover (*k*_cat_) and >5-fold increase in *K*_M_, resulting in >3000-fold reduction in catalytic efficiency (*k*_cat_/*K*_M_) relative to wtOXA-48. Similarly, introduction of V120G into wtOXA-48 caused a 3-fold decrease in *k*_cat_ and a 3-fold increase in *K*_M_, resulting in a 9-fold reduction in *k*_cat_/*K*_M_. These reductions in catalytic parameters are consistent with the observed decreases in PIP *IC_50_* values (Tab. 1). Importantly, the trade-off–mitigating variant Q4:V120G displayed a 3-fold reduction in *k*_cat_ but a >5-fold improvement in *K*_M_ relative to Q4. Thus, the enhancement in Q4:V120G-mediated PIP resistance is likely driven by an improved overall *K*_M_.

**Tab. 1:** Biochemical characterization of wtOXA-48, Q4, and their respective V120G variants.

| OXA-48<br>variants | $IC_{50}$<br>(mg/L) | Steady-state parameters | | | $T_M$<br>(°C) |
| --- | --- | --- | --- | --- | --- |
| | | $k_{\text{cat}}$ (s<sup>-1</sup>) | $K_M$ (μM) | $k_{\text{cat}}/K_M$ (M<sup>-1</sup> s<sup>-1</sup>) | |
| wtOXA-48 | 16.9 ± 0.6 | 150 ± 20 | 125 ± 80 | 1.2x10<sup>6</sup> | 53.0 |
| wtOXA-48:V120G | 1.2 ± 0.1 | 50 ± 2 | 370 ± 35 | 1.3x10<sup>5</sup> | 51.4 |
| Q4 | 0.6 ± 0.1 | 0.47 ± 0.10 | >600 | 3.6x10<sup>2</sup> | 45.0 |
| Q4:V120G | 2.6 ± 0.2 | 0.14 ± 0.01 | 200 ± 25 | 6.9x10<sup>2</sup> | 46.8 |

To investigate potential non-catalytic compensatory effects of V120G, we analyzed enzyme stability by determining the thermal denaturation midpoint (*T_M_*) for wtOXA-48, Q4, and their corresponding V120G variants (Fig. 2d, Tab. 1). Q4 displayed reduced stability compared to wtOXA-48, with an 8°C decrease in *T_M_*. This is consistent with previous results showing that F72L, in several variants across the Q4 evolutionary trajectory, lowers stability by 6°C when introduced into wtOXA-48^36^. Furthermore, V120G decreased stability by 1.5°C in the wtOXA-48 background but increased stability by 1.8°C in Q4, suggesting a modest restabilizing effect of V120G specifically in the Q4 background.

Overall, co-selection yielded a Q4:V120G variant that partially restored PIP resistance, thereby mitigating the initial collateral sensitivity trade-off between PIP and CAZ (Tab. S3). Incorporation of V120G into the adaptive landscape revealed that this mutation had minimal impact on CAZ resistance, whereas its effect on PIP resistance depended strongly on the presence or absence of F72L (Fig. 2b, Fig. S1-S2, Tab. S4-S5). Our biochemical investigation indicates that the PIP trade-off introduced by Q4 mutations arises from a combination of reduced *k*_cat_ and *K*_M_, while PIP mitigation in Q4:V120G is seemingly driven by improvements in *K*_M_ (Fig. 2c, Tab. 1).

### Systematic substitutions at position 120 revealed trade-off mitigation beyond glycine

V120 is crucial for OXA-48’s activity by shaping the deacylation water channel and assisting the hydrophobic environment around the catalytic base^27,37,38^. To assess the role of position 120 in balancing PIP and CAZ resistance, we systematically substituted V120 with all naturally occurring amino acids in Q4 by site-directed mutagenesis and determined *IC_50_* values for CAZ and PIP (Fig. S3a, Tab. S6). Specifically, we asked whether other amino acid substitutions at position 120 exist which could mitigate the observed PIP trade-off.

Among the Q4:V120X variants, three substitutions (V120I, V120W, and V120D) reduced resistance to both antibiotics relative to Q4. Two variants (V120N and V120E) were deemed neutral, showing no gain in PIP *IC_50_* and a <2-fold decrease in CAZ resistance. V120M, V120F, and V120C modestly increased PIP *IC_50_* (1.5-2-fold) but at the cost of reducing CAZ *IC_50_* values (2-6-fold), thereby still demonstrating collateral sensitivity. V120Y produced the strongest gain in PIP resistance (15-fold) but concurrently led to a detrimental decrease in CAZ resistance (73-fold).

In contrast, several substitutions displayed trade-off–mitigating behavior in Q4. Three non-polar, hydrophobic substitutions (V120L, V120A, and V120P), in addition to the polar V120Q, increased PIP *IC_50_* by 2 to 4-fold. In particular, V120L and V120A achieved 4-fold increases in PIP *IC_50_* without substantial loss in CAZ *IC_50_* (<2-fold), similar to V120G. Likewise, all positively charged (V120R, V120H, and V120K) and two polar substitutions (V120S and V120T) either maintained or increased PIP *IC*_50_ values up to 4-fold, while simultaneously improving CAZ resistance by up to 2-fold.

Next, we asked whether introducing the same amino acid substitutions would have similar effects in the wtOXA-48 background. None of the substitutions at position 120 altered CAZ resistance in wtOXA-48 (Fig. S3b, Tab. S7). In addition, analysis of PIP *IC*_50_ values in wtOXA-48:V120X variants showed that most substitutions were highly detrimental to PIP resistance, resulting in 3- to 71-fold reductions relative to wtOXA-48. In fact, only aromatic substitutions (V120W, V120F, and V120Y) maintained PIP *IC*_50_ values similar to wtOXA-48 (<2-fold change). This stands in strong contrast to Q4:V120X variants, in which the same substitutions produced diverse effects on PIP resistance (3-fold decrease for V120W, 2-fold increase for V120F, and 15-fold increase for V120Y).

Together, these results indicate that the PIP-trade-off-mitigating effect in Q4 is not unique to glycine and can be achieved by a variety of other amino acids, including certain hydrophobic (V120A, V120P, and V120L), polar (V120S and V120T), and positively charged (V120R, V120H, and V120K) residues (Fig. S3a, Tab. S6). Moreover, the strikingly different effects of the same substitutions in wtOXA-48 and Q4 suggest that mutations accumulated in Q4 reshape the local structural and chemical environment around position 120, thereby modulating both PIP and CAZ resistance (Fig. S3b, Tab. S6-S7).

### Changes in Ω-loop dynamics accompany collateral sensitivity and its mitigation

During the evolution of wtOXA-48 to Q4, the Ω-loop experiences a substantial increase in flexibility and a shift in its orientation, a process mainly driven by F72L^36^ (Fig. 3a). We therefore hypothesized that changes in Ω-loop conformational freedom and orientation may be related to the observed PIP trade-off and its mitigation. To test this, we crystallized and soaked the endpoint mutant from our evolutionary trajectory, Q5 (A33V/K51E/F72L/S212A/T213A)^36^, as it displays a resistance profile highly similar to that of Q4 (Tab. S1). The structure of Q5 in complex with PIP (Q5-PIP, PDB: 9TAZ) was solved with one protein molecule (chain A) in the asymmetric unit at a resolution of 1.5 Å (Tab. S8, Fig. S4). PIP was captured in an acyl-enzyme state, with two additional PIP molecules coordinated outside of the active site. In Q5-PIP, residues V153-L158 in the Ω-loop could not be rebuilt due to a lack of electron density, indicating a highly mobile state of this loop region (Fig. 3b). In addition, a chloride ion was observed in close proximity to PIP and the conserved residue R250, which anchors the C3-carboxyl group of β-lactams *via* electrostatic interactions^30,37–40^ (Fig. 3b-c, Fig. S4).

**Fig. 3:**
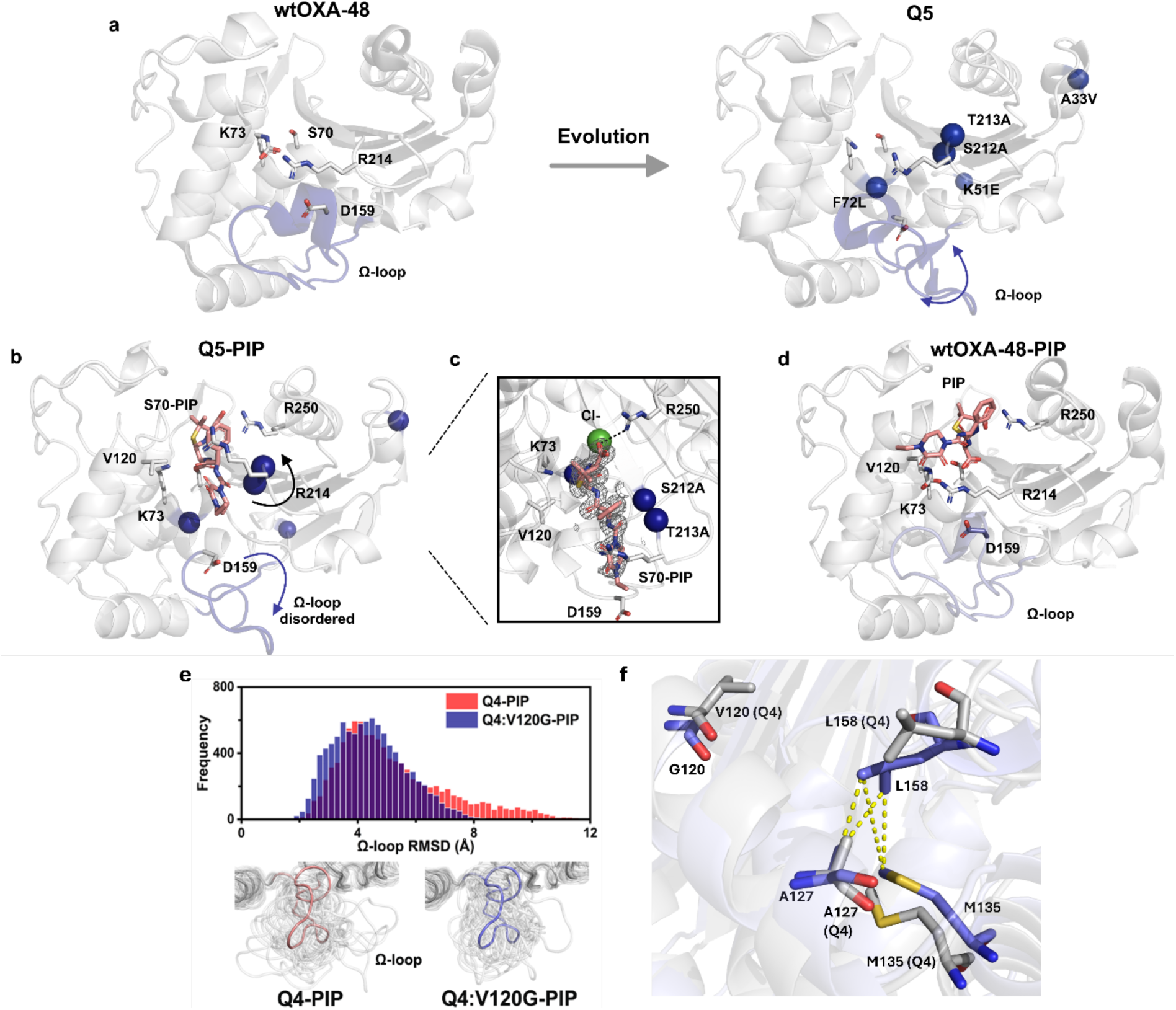
Changes in Ω-loop orientation and its conformational space during evolution. **a.** In wtOXA-48 (left, PDB: 4S2P^41^), the Ω-loop is stabilized by a salt bridge between R214 and D159 in the loop region. Residues S70, K73, D159, and R214 are highlighted. In Q5 apo (right, PDB:8PEB^36^), the Ω-loop is displaced by a shift in orientation due to increased flexibility. Q5 mutations are shown with blue spheres. **b.** The R214 backbone and side chain in Q5-PIP (PDB: 9TAZ) adopt an alternative orientation compared to wtOXA-48 (Cα shifted by 3 Å), thereby avoiding a clash between R214 and PIP. **c.** A chloride ion is located in the active site of Q5-PIP, in close proximity to R250 and the C3-carboxyl group of piperacillin that can form a salt bridge. Electron density of piperacillin is shown based on the 2fo-fc map contoured at 1 σ. **d.** Docking of wtOXA-48 with piperacillin shows that the substrate adopts a different orientation compared with Q5-PIP. **e**. Cα RMSD histograms (0.2 Å bin width, top) and ensembles (bottom) of the Ω-loop (Y144 to R163) obtained from MD simulations. Q4:V120G’s Ω-loop samples a smaller conformational space compared to Q4. **f.** V120G facilitates an alternative conformation of L158 in Q4:V120G, promoting more frequent interactions with A127 and M135 compared to Q4 (see Fig. S6). Q4 residues are shown in gray, while Q4:V120G residues are shown in blue.

While we repeatedly attempted to obtain crystal structures of wtOXA-48 in complex with PIP, the obtained datasets showed weak electron density for PIP in the active site, preventing reliable modeling (Fig. S5a). Therefore, these datasets were complemented with molecular docking. The docking procedure was validated by docking hydrolyzed and non-hydrolyzed PIP into Q5, which reproduced the PIP conformation observed in the Q5-PIP structure (Fig. S5b-c, Tab S9). We then calculated the center of mass based on the PIP position of the soaked wtOXA-48 crystals, which served as a starting point for molecular docking in wtOXA-48. The resulting docked wtOXA-48-PIP structure matched well with the placement of PIP based on the partial electron density (Fig. S5d-e, Tab S9).

For structural comparison, we aligned Q5-PIP with the apo structures of wtOXA-48 (PDB ID: 4S2P^41^), Q5 (PDB ID: 8PEB^36^), and our docked wtOXA-48-PIP structure. Our analysis revealed a steric clash between PIP in Q5 and the side chain of R214 in wtOXA-48 (Fig. 3a,b). In fact, PIP binding in Q5 appears to displace R214 by ∼3 Å (Cα movement) relative to wtOXA-48. R214 forms strong electrostatic interactions with D159 in the Ω-loop, keeping the active site in a closed state even in acylated forms in wtOXA-48^42–44^ (Fig. 3a). This interaction weakens during evolution from wtOXA-48 to Q4/Q5, allowing the Ω-loop to sample a different conformation and a broader conformational space. Analyzing PIP binding in wtOXA-48 and Q5 further uncovered substantially different substrate binding poses due to the evolutionary remodeling of the Ω-loop (Fig. 3d). Consequently, combined results from X-ray crystallography and molecular docking indicate that evolutionary changes in Ω-loop dynamics give rise to alternative substrate binding poses, which likely underlie the observed collateral sensitivity (Fig. 3a-d, Fig. S5, Tab. S8-S9).

Next, we aimed to investigate the potential structural role of V120G on the Ω-loop. For this, molecular dynamics (MD) simulations were performed on the PIP acyl-enzymes structures of Q4 and Q4:V120G. Based on 32 independent MD trajectories each, the Ω-loop (Y144-R163) in Q4:V120G sampled a substantially smaller conformational space (4.4 Å) compared to Q4 (5.2 Å), indicating decreased conformational freedom (Fig. 3e). In fact, the simulations indicate that V120G facilitates a conformational change in L158, allowing L158 to form more stable hydrophobic interactions with A127 and M135 (Fig. 3f, Fig. S6), potentially contributing to stabilization of the Ω-loop after the introduction of F72L.

In summary, increased Ω-loop flexibility during CAZ evolution of OXA-48 to Q4/Q5 facilitates CAZ accommodation but likely results in less productive PIP-binding poses, potentially causing the observed strong trade-off. V120G, which mitigates the PIP trade-off in Q4, appears to stabilize the Ω-loop by strengthening hydrophobic interactions.

## DISCUSSION

Collateral sensitivity is being explored as an evolutionary constraint that can be exploited to design efficient antimicrobial therapies^1–4^. However, its success hinges on evolutionarily robust collateral networks, and numerous studies have shown that collateral effects are often transient and context dependent, allowing rapid mitigation or even inversion of collateral responses^1–3,10,16,21,22^. Here, we provide a mechanistic understanding of the evolutionary stability of collateral networks at the level of a single resistance determinant, using the β-lactamase OXA-48 as a model system. We show that during evolution, increased Ω-loop flexibility likely promotes sampling of imperfect substrate binding poses, which can give rise to collateral sensitivity. Under co-selective pressure, however, single substitutions can evade these constraints by readjusting the conformational freedom of this loop (Fig. S3, Tab. S6-S7). These findings illustrate that collateral networks are not inherently robust and can be eroded by adaptive processes. Similar events have been observed on the population level of *E. coli* and *Pseudomonas aeruginosa*, where adaptive evolution under alternating or combined drug pressures led to the collapse of collateral constraints^10,16,45–47^. Notably, in our study, mitigation occurred only in the Q4 background, not in wtOXA-48, demonstrating the importance of the evolutionary starting point in overcoming such trade-offs.

Similar to OXA-48, comparable collateral sensitivity trade-offs have been reported during the evolution of OXA-48-like variants as well as other serine β-lactamases (e.g., TEM, KPC, and CTX-M variants)^23,25,30,32,35,39,42,48,49^. In these enzymes, the Ω-loop is a well-established evolutionary target that helps determine substrate specificity. Increased flexibility within this loop region is commonly associated with evolution for cephalosporin resistance and often occurs at the expense of penicillinase or carbapenemase activity^42,49–55^. In Q4, F72L disrupts an aromatic pocket (including Y144 and F156) within the active site^36^. The resulting increase in Ω-loop flexibility likely facilitates CAZ accommodation but impairs PIP hydrolysis due to imperfect substrate binding (Fig. 3). V120G appears to mitigate this trade-off by rigidifying the Ω-loop, consistent with a general improvement in *K*_M_ (Tab. 1, Fig. 3e-f). The changes in flexibility are further supported by thermostability data, which show an 8 °C decrease in *T*_M_ in Q4 relative to wtOXA-48 and a 1.8 °C increase in Q4:V120G compared to Q4 (Tab. 1, Fig. 2d). These observations provide insight into the molecular principles governing collateral responses in β-lactamases and how evolutionary trade-offs emerge and are reshaped during adaptation.

Our results provide a mechanistic perspective on the evolutionary robustness of collateral sensitivity at the level of a single resistance enzyme. Using OXA-48 as a model system, we demonstrate that collateral sensitivity can arise from changes in loop dynamics and that co-selection with antibiotics exhibiting collateral sensitivity can mitigate these effects. These results underscore the importance of a detailed understanding of the molecular mechanisms shaping collateral sensitivity to distinguish robust from transient trade-offs and to design treatment strategies that preserve exploitable sensitivities over time.

## METHODS

### General material

Lysogeny broth (LB), LB agar, chloramphenicol, ceftazidime, and piperacillin were purchased from Sigma-Aldrich (Saint-Louis, MO, USA). All enzymes used for cloning were purchased from Thermo Fisher Scientific (Waltham, MA, USA). The cloning strain *E. coli* E. cloni 10G was purchased from Lucigen. All primers were purchased from IDT (Integrated DNA Technologies, IA, USA; Tab. S10). All kinetic and *IC_50_* measurements were calculated in GraphPad Prism v. 10.6.0 (GraphPad Software, CA, USA).

### Directed evolution, cloning, and co-selection

Mutational libraries were constructed for wtOXA-48 and the Q4 variant using error-prone polymerase chain reaction to introduce random mutations into the *bla*_OXA-48_ gene. The *bla*_OXA-48_ gene was located on a low to medium copy plasmid (origin pA15)^48^. PCR reactions were performed with 10 ng DNA template, 10 mM dNTPs, 25 mM MgCl_2_, 10 µM preOXA forward/reverse primers (Tab. S10), GoTaq G2 polymerase (Promega, WA, USA), and either 500 µM 8-oxo-dGTP or 10 µM dPTP (Jena Bioscience, Germany). PCR products were purified (QIAquick PCR purification kit, Qiagen, Germany) and digested with *DpnI* for 1 h at 37 ℃, 400 rpm to remove template DNA. A second PCR was carried out as described previously, using 5 ng of each purified product, but without mutagenic nucleotides. PCR products were cleaned again and digested with *NcoI* and *XhoI* for 1 h at 37 ℃, 400 rpm to create sticky ends, then ligated into the vector backbone using T4 DNA ligase for 2 h at 22.5 ℃. The ligated product was purified using the MicroElute kit (Omega Bio-Tek, GA, USA) and transformed into E. cloni 10G electrocompetent cells (Lucigen, WI, USA). To ensure sufficient mutational depth, libraries were plated on LB agar with 25 mg/L chloramphenicol to confirm a yield of >5000 colonies per library. Co-selection of established mutational libraries was performed on LB agar plates containing 0.15 or 0.5 mg/L CAZ in combination with 0.5, 1, 2, 4, or 8 mg/L PIP (Tab. S2). Plates were incubated at 37 ℃ for 20 h, and colonies surviving under the highest antibiotic concentrations were recovered. Potential mutations in the *bla*_OXA-48_ gene were identified by Sanger sequencing using preOXA primers (Tab. S10). Additionally, the wtOXA-48:V120G and Q4:V120G variants were constructed using Phusion polymerase and no mutagenic nucleotides with the OXA_V120G forward/reverse primers (Tab. S10). All variants were subcloned into isogenic vector backbones and *E. coli* E. cloni strains before the *IC*_50_ determination.

### Half-maximal inhibitory concentration (*IC_50_*) susceptibility testing

Susceptibility testing was performed by microbroth dilution in 384-well plates (Thermo Fisher Scientific) using two-fold serial dilution of either PIP or CAZ, as previously described^29,48^. In summary, LB broth was inoculated with 10^5^ - 10^6^ CFU/mL and incubated at 37 °C for 20 h. Bacterial growth was measured at OD_600_ using a microplate reader (BioTek Epoch 2, Agilent Technologies, CA, USA). Dose-response curves and *IC_50_* values were calculated in GraphPad Prism v. 10.6.0.

### Protein expression and purification

Protein expression was performed as described previously^36^. In short, *E. coli* BL21-AI cells carrying wtOXA-48/V120G (MP21-1/MP29-26), Q4/V120G (MP12-48/MP29-44), Q5 (MP21-73), in a pDEST-17 expression vector were incubated overnight in LB supplemented with 100 mg/L ampicillin. Overnight cultures were used to inoculate terrific broth containing 100 mg/L ampicillin and grown at 30 °C, 220 rpm until an OD_600_ of 0.4 was reached. Expression was induced by adding 0.2% L-arabinose and 0.1 mM IPTG, followed by incubation at 15 °C, 225 rpm overnight. Cells were collected by centrifugation and resuspended in buffer AL (50 mM HEPES pH 7.2, 50 mM K_2_SO_4_, 0.1% Triton, 1x cOmplete protease inhibitor). The cell suspension was sonicated on ice for 1 h, and cell debris was removed by centrifugation at 15,000 rpm for 10 min at 4°C. The supernatant was purified using HisPur Ni-NTA spin columns (Thermo Fisher Scientific). Bound proteins were washed with buffers containing 50 mM HEPES (pH 7.2), 50 mM K_2_SO_4_, and 25 mM imidazole, then subsequently eluted with 250 mM imidazole in the same buffer. Eluted samples were concentrated using Amicon Ultra (Merck Millipore, Burlington, MA, USA) at 4°C, 15,000 rpm. Final protein concentrations were quantified by measuring *OD*_280_ and subsequently calculated using the respective extinction coefficients.

### Steady-state enzyme kinetics

Michaelis-Menten catalytic parameters (*k*cat, *K*_M_, and *k*_cat_/*K*_M_) for OXA-48 variants were determined under steady-state conditions for piperacillin (Δε = −820 M−1 cm−1, 235 nm) by measuring initial reaction rates in a spectrophotometer (BioTek Epoch 2, Agilent Technologies). Final enzyme concentrations were 1 nM for wtOXA-48 and wtOXA-48:V120G, and 250 nM for Q4 and Q4:V120G. Measurements were conducted in at least duplicate in UV-transparent 96-well plates (Sigma-Aldrich) at a final assay volume of 100 µL in buffer containing 0.1 M KH_2_PO_4_ (Sigma-Aldrich), 0.1 M K_2_HPO_4_ (Sigma-Aldrich), and 50 mM NaHCO_3_ (Sigma-Aldrich). Substrate concentrations ranged from 10 µM to 600 µM for all variants and up to 1250 µM for wtOXA-48. All measurements were performed at 25 °C, and kinetic parameters were obtained by nonlinear regression fitted to the Michaelis-Menten equation in GraphPad Prism v10.6.0 (GraphPad Software).

### Thermostability assay

The thermal stability of purified 6x His-tagged OXA-48 variants containing TEV protease cleavage site was measured as previously described^29,36^. Enzymes were diluted to a final concentration of 0.2 mg/ml in 50 mM HEPES (pH 7.5, VWR) with 50 mM potassium sulfate (Honeywell, NC, USA). Each sample was mixed with 5x SYPRO Orange dye (Sigma-Aldrich) and exposed to a controlled temperature gradient from 25 °C to 70 °C in an MJ Mini^TM^ Gradient Thermal Cycler (Bio-Rad, CA, USA) at a heating rate of 1 °C/min. Fluorescence changes were monitored in triplicates, and the thermal denaturation midpoint (*T_M_*) was determined.

### Site-directed mutagenesis

Site-directed mutagenesis at position V120 of the Q4 variant of OXA-48 was achieved by PCR with a primer incorporating the degenerate codon NNS that encodes all 20 amino acids, creating a mutational library of all variants in this specific site. PCR reactions were performed using 10 ng DNA template (pUNe-4^36^), 10 mM dNTPs, 10 µM forward/reverse primers (Tab. S10), Phusion polymerase (Thermo Fisher) in 5x Phusion HF buffer. PCR products were cleaned (QIAquick PCR purification kit, Qiagen) and digested with *DpnI* and *LguI* for 1 h at 37 ℃, 400 rpm to remove template DNA and primers. Self-ligation of the insert was performed using T4 DNA ligase for 2 h at 22.5 ℃ before transformation into E. cloni 10G electrocompetent cells (Lucigen) and incubated overnight on LB agar containing 25 mg/L chloramphenicol at 37 ℃. Single clones from the resulting library were selected and screened on agar plates supplemented with different concentrations of CAZ and PIP, and phenotypically distinct clones were subjected to Sanger sequencing. Using this approach, 12 of the 19 variants were successfully isolated. To obtain the remaining amino acid substitutions and complete the full set of natural variants, site-specific primers were designed and used for targeted mutagenesis (Tab. S10).

### Crystallization and structural refinement

Q5-PIP was crystallized as previously described^36^. Crystals were soaked for 15 min, cryoprotected with 15% ethylene glycol (Sigma-Aldrich) in the reservoir solution, and subsequently frozen in liquid nitrogen. Diffraction data were collected on ID23-2 (Q5-PIP)^56^ at the European Synchrotron Radiation Facility (ESRF), Grenoble, France, at 100 K, wavelength 0.873 Å, and the diffraction images were indexed and integrated using XDS^57^. For data scaling, AIMLESS^58^ was used, aiming for overall high completeness and CC1/2 > 0.5, and a mean intensity above 1.0 in the outer resolution shell (Tab. S8). Molecular replacement was performed using chain A from PDB ID 6Q5F^59^ and the program PHENIX v. 1.12^60^. Parts of the model were manually rebuilt using Coot^61^. Average structure and ensemble refinement were performed using PHENIX v. 1.12. PyMOL 3.1.3 was used for illustrations (Schrödinger, NY, USA).

### Molecular docking

Molecular docking studies were performed to predict possible binding modes of hydrolyzed and non-hydrolyzed PIP towards wtOXA-48 and Q5-PIP. Structures used for molecular docking were obtained from crystal structures of wtOXA-48 (PDB:4S2P^41^) and Q5-PIP (PDB:9TAZ). Protein structure preparation was carried out with the Protein Preparation Wizard^62^ in Schrödinger Release 2020, at pH 7.0 using the OPLS3e force field. The docking grid was generated with the Receptor Grid Generation module in Glide. Ligand preparation for piperacillin and its hydrolyzed form was performed using the LigPrep module (Schrödinger) at pH 7.0, generating low-energy three-dimensional conformations. Flexible molecular docking was performed in Glide with Standard Precision (SP) mode^63,64^ (Schrödinger), and the three best-ranked docking poses were retained for subsequent analysis.

### MD simulations

Models of the Q4 and Q4:V120G piperacillin (PIP) acyl-enzyme complexes were generated based on the Q5-PIP crystal structure (PDB ID: 9TAZ). Ω-loop residues 151 to 161 were predicted using Modeller^65^, and the carbamylation on Lys73 (KCX) and the deacylating water were added manually. Water molecules from the structure were retained, and ionizable residues were in their standard protonation states, consistent with p*K*_a_ predictions at pH 7.5 from PropKa3.1^66^, with all histidine residues singly protonated on NE2 (as predicted by reduce from AmberTools^67^). The complexes were solvated in a rectangular box of TIP3P water^68^, neutralized by replacing 3 bulk waters with Na^+^. The Amber ff14SB force field^69^ was used for the protein, with consistent KCX parameters from previous work^70^ and for the PIP-acetylated serine (partial charges calculated from restrained electrostatic potential fitting with the RED Server^71^, and bonding and Van der Waals parameters described by the GAFF2 force field).

MD simulations were performed analogously to our previous work^36^, using Amber20^67^ In short, after minimization and brief heating (50 to 300 K in 20 ps) with restraints to maintain productive KCX and deacylating water positions, 40 ns MD simulation in the NPT ensemble at 300 K and 1 bar was carried out (Langevin dynamics with 2 fs time-step with SHAKE applied, and Berendsen barostat). 32 independent simulations were conducted per system (1.28 μs total sampling). The initial 10 ns were discarded (as unrestrained equilibration). Analysis was conducted using CPPTRAJ^72^ on frames saved every 100 ps from 10-40 ns MD, resulting in 9600 frames per system. C*α* Root mean square deviations (RMSD) of the Ω-loop were calculated by aligning trajectories on all C*α* atoms excluding those in the Ω-loop.

## DATA AVAILABILITY

The cryogenic crystal structure was deposited to the Protein Data Bank under the PDB ID: 9TAZ (Q5-PIP). Docked structures will be made available on GitHub upon publication. Starting structures, topology files, and input files for MD simulation, as well as the required parameter files for the PIP-acylated serine will be made available upon publication.

## ACKNOWLEDGEMENTS

DS and CF were supported by the Centre for New Antibacterial Strategies (CANS) at UiT - The Arctic University of Norway. CF thanks the Norwegian Monitoring Systems for Antibiotic Resistance in Microbes (NORM), the Federation of European Biochemical Societies, the Federation of European Microbiological Societies, and the Northern Norway Regional Health Authority (HNF1722-24). DW and MWvdK thank the GW4 BIOMED2 DTP for DW’s studentship, grant MR/W006308/1 awarded to the Universities of Bath, Bristol, Cardiff, and Exeter from the Medical Research Council (MRC)/UKRI. We acknowledge ESRF for the provision of synchrotron-radiation facilities under proposal number MX-2332 and thank the staff of the ESRF and EMBL Grenoble for assistance and support in using beamline ID23-2. MD simulations were conducted using the computational facilities of the Advanced Computing Research Centre, University of Bristol. KVG gratefully acknowledges the Swedish Research Council (2024-05496) and the Uppsala Antibiotic Center for financial support.

## AUTHOR CONTRIBUTIONS

DS and CF performed and analyzed the biochemical assays. CF crystallized and solved structures. KVG performed the molecular docking experiments. KB analyzed and visualized data. DW and MWvdK performed and analyzed MD simulations. CF designed the study. DS and CF wrote the manuscript with input from all coauthors.

